# Strand Displacement Activity of Mimiviral Polymerase X Enables Rapid Detection of Sequence-Specific DNA Targets

**DOI:** 10.64898/2026.06.10.731134

**Authors:** Apoorva S Raman, Shailesh B. Lad, Soumyadeep Mandal, Debjani Paul, Kiran Kondabagil

## Abstract

Mimiviral polymerase X, mvPolX, is a repair polymerase that is involved in base excision repair (BER) and carries out the gap-filling function in double-stranded DNA (dsDNA). We demonstrate a sensitive and sequence-specific DNA detection method using this polymerase. mvPolX begins polymerizing DNA from the 3’ end of a gap, displacing the downstream nucleotides without exonuclease activity. Our detection method is built on this activity of mvPolX. We designed a probe molecule consisting of a partial dsDNA with a 3’ over-hang region complementary to the target DNA to be detected. The probe has a fluorophore-quencher (FAM-BHQ1) tag to facilitate detection upon strand removal. Binding of the probe to the complementary target forms a dsDNA with a single nucleotide gap in one strand. mvPolX binds this gap region and begins polymerisation eventually displacing the quencher strand leading to an increase in fluorescence. Proof-of-concept has been established using a synthetic 19 bp target DNA sequence. The method is specific and did not show any strand displacement when a single or double mismatched nucleotide at the 3’ end of the target DNA was used. To demonstrate this molecular assay, we used M13 phage as our target. Asymmetric PCR (aPCR) was used to obtain single-stranded target DNA (158 bases) from M13 genomic DNA, which was directly used in the assay as target. The combination of aPCR and mvPolX assay can detect as low as 10 copies of genomic DNA. The enzymatic reaction is fast, requiring only 15 min of incubation with mvPolX at 30 °C. We have further demonstrated the efficiency of the assay in presence of multiple non-target DNA by detecting the target DNA from M13 phage spiked lakewater samples.

## 1 Introduction

While viruses range in size from 20-200 nm, giant viruses are larger than 300 nm.^1,2^ Their genome sizes range from a few hundred kilobases to megabases.^2,3^ Beyond performing simple proliferative functions, giant viruses also code for proteins that can repair their own DNA. For example, their Base Excision Repair (BER) mechanism corrects a damaged base in the viral DNA. X DNA polymerases (PolXs) are a family of repair polymerases that work in conjugation with other enzymes to perform BER.^4–6^ These polymerases can identify and carry out the function of gap filling during DNA repair.^7,8^ Compared to the replicative polymerases, PolXs are smaller in size and exhibit a lower fidelity of DNA polymerization.^9^

A mimiviral repair polymerase X, mvPolX, was recently identified and characterized.^10^ mvPolX is a monomeric protein of 42 kDa. Its active site consists of three aspartate residues at the 201^st^, 203^rd^ and 269^th^ positions of the amino acid chain.^9–11^ mvPolX can extend a DNA strand from its 3’ OH end. Most importantly, the polymerisation can take place successfully even if there is a blocking primer present downstream with a single or a five-nucleotide gap. It should be noted that mvPolX has no exonuclease activity owing to its lack of proofreading.^10^

Here, we exploited the polymerization-dependent strand displacement activity of mvPolX to design and demonstrate a sequence-specific DNA detection method. We have designed a probe molecule by exploiting the fact that mvPolX carries out polymerization only in presence of a target DNA molecule that has a perfect sequence match. The probe molecule is a partially double-stranded DNA with a fluorophore (FAM) tagged to the longer strand (‘target-binding strand’) and a quencher (BHQ1) tagged to the shorter strand (Fig. 1A). The single-stranded overhang region of the probe is designed to be complementary to the target DNA sequence. When the target sequence binds to the single-stranded region of the probe molecule, the complete structure resembles that of a double-stranded DNA, but with a one-nucleotide gap in one of the strands. mvPolX binds to this gap and starts polymerizing DNA from the 3’ OH end of the target sequence. As it continues polymerization, mvPolX displaces the shorter strand of the probe molecule. Since the shorter strand containing BHQ1 is removed, there is an increase in the FAM fluorescence, indicating the presence of the target molecule (Fig. 1B). The mvPolX assay takes place at 30°C and gets completed in 15 min, leading to sequence-specific detection of DNA targets. We showed the proof of concept of this assay using a synthetic target as well as a 158-base sequence from the M13 phage. We demonstrated the specificity of this assay using a non-complementary synthetic target as well as with synthetic targets with single and double mismatches. We believe with additional benchmarking, this molecular assay can be a suitable candidate for use in DNA-based diagnostics.

**Figure 1.**
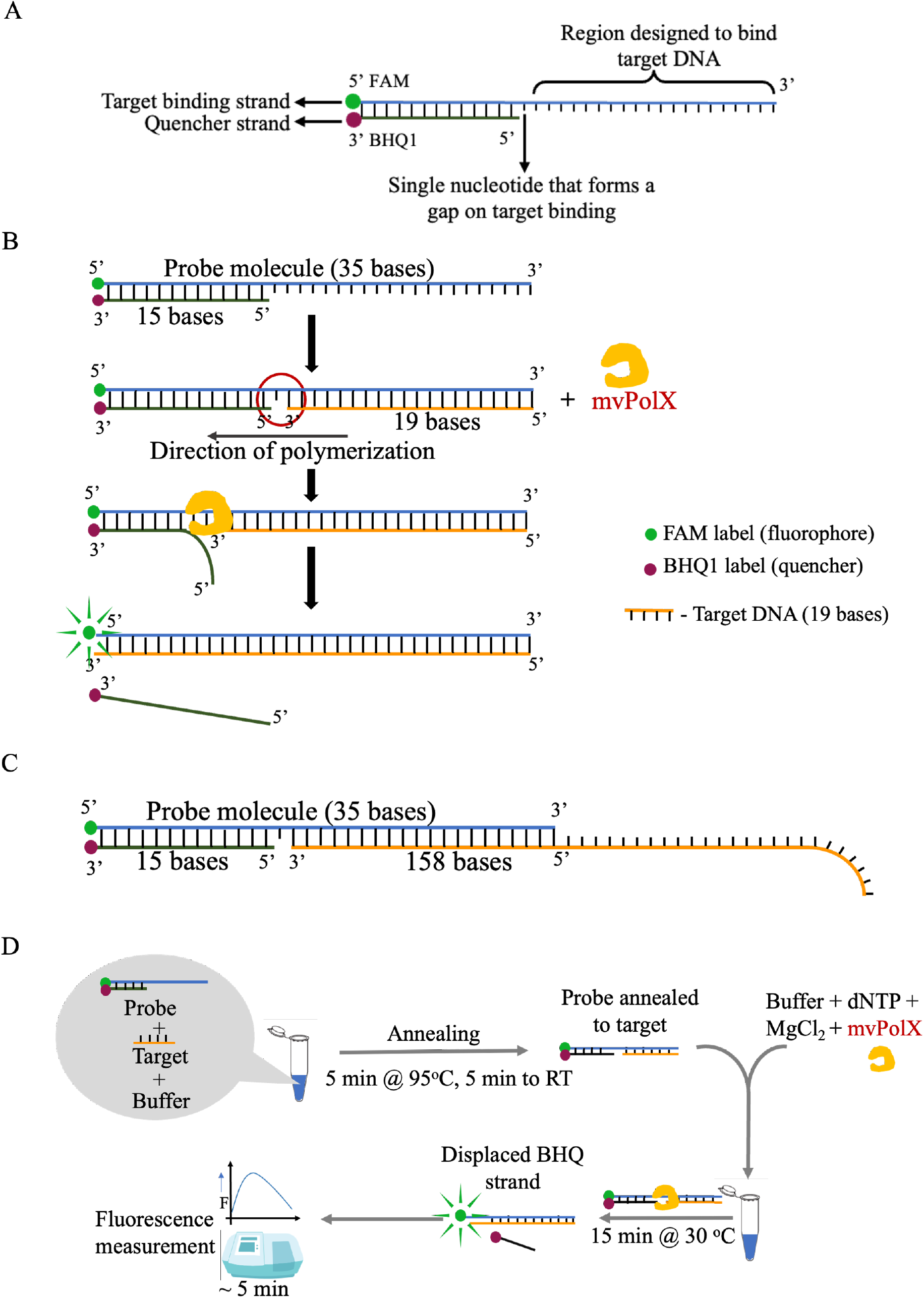
mvPolX assay principle. (A) Schematic showing the probe design (B) Schematic diagram showing the principle of the mvPolX assay; (C) Schematic showing single-stranded DNA bound to probe; (D) Schematic representation of the assay protocol

## 2 Materials and methods

### 2.1 Sequences of the probe molecule and synthetic DNA targets

The sequences of the FAM-labelled and BHQ1 labelled DNA strands in the probe molecule are 5’-[FAM]TCCTACCGTGCCTACTGAACGGGTATTAAACCAAG-3’ and 5’-GTAGGCACGGTAGGA[BHQ1]-3’ respectively. We designed a 19 bp synthetic target oligonucleotide with the sequence 5’-CTTGGTTTAATACCCGTTC-3’. The probe and the synthetic target oligonucleotides were synthesized by Eurogentec, Belgium. A non-complementary synthetic construct with the sequence 5’-GACCCCAAAATCAGCGAAAT-3’ was used to check specificity of the assay. Further, two target oligonucleotides with sequences 5’-CTTGGTTTAATACCCGTTA-3’ and 5’-CTTGGTTTAATACCCGTCA-3’ were synthesized with a single base and a double base mismatch respectively. The non-complementary synthetic target and the target oligonucleotides with single and double mismatches were synthesized from Sigma-Aldrich, India.

### 2.2 Culture of M13 phage

*E. coli* XL1-Blue strain was used as the host organism to culture the M13 phage. *E. coli* was cultured overnight at 37°C and 150 rpm. 200 *µ*l of diluted (by a factor of 10^5^) mid-log phase *E. coli* culture was mixed with 200 *µ*l of M13 phage suspension (6 × 10^2^ phage particles/ml) in sodium chloride - magnesium sulphate (SM) buffer to maintain a multiplicity of infection (MOI) of 0.5. This suspension was mixed with melted top agar (0.4 %) and was poured on to a LB agar plate and spread evenly. The plate was allowed to dry at room temperature before shifting to 37°C for overnight incubation.^12^ This method was used for preparation of phage suspensions and estimation of PFU/ml from liquid cultures.

For isolation of phage genomic DNA, 200 *µ*l of diluted overnight *E. coli* culture was added to 300 *µ*l of M13 phage solution and made up to a total volume of 1 ml with Luria-Bertani broth. The MOI was maintained at 0.5. The suspension was incubated at 37°C for 2 h and then added to 20 ml Luria broth. This suspension was incubated at 95 rpm for 7 h at 37°C before harvesting for isolation of genomic DNA.^12^

### 2.3 Isolation of M13 genomic DNA

For extraction of M13 genomic DNA, the liquid culture was centrifuged at 3000 *g* for 10 min to separate the bacteria from phage particles. The supernatant was collected carefully without disturbing the pellet. To this supernatant, NaCl was added to a final concentration of 1M and incubated the on ice for 1 h. Next, PEG 8000 solution was added to a final concentration of 10% w/v and the solution was kept at 4°C overnight to precipitate the phage particles. Then the solution was centrifuged at 21000 *g* for 20 min. The supernatant was discarded and the pellet containing phages was resuspended in 200 *µ*l of sterile Milli-Q water. The phage suspension was treated with DNase I (0.5 U) and RNase (final concentration 20 μg/ml) at 37°C for 1 h. It was next treated with Proteinase K (final concentration 20 μg/ml) in presence of sodium dodecyl sulphate (SDS) (final concentration 0.5%) and ethylenediaminetetraacetic acid (EDTA) (final concentration of 20 mM) for 2 h at 60°C. Once cooled to room temperature, the solution was extracted with an equal volume of a phenol: chloroform: isoamyl alcohol mixture (25:24:1). The extraction was repeated twice, followed by a final extraction with only chloroform. Next, 1/10^th^ of the volume of 3M Sodium acetate was added, followed by addition of 2.5 times the volume of chilled 100% ethanol. The solution was kept at −20°C overnight to precipitate the DNA, followed by centrifugation at 21000 *g* for 20 min.^12^ The DNA pellet was washed with 70% ethanol, dried and redissolved in 50 *µ*l of nuclease-free water. The concentration of the purified DNA was measured using the NanoDrop Lite Spectrophotometer (Thermo Scientific, India). The extracted M13 genomic DNA was directly used as template for asymmetric PCR preceding the mvPolX assay.

### 2.4 Amplification of the target gene by asymmetric PCR

A 158 bp sequence of the gene 1 region was amplified using the isolated M13 phage genome as the template. The forward and reverse primer sequences were 5’-GTCGGGAGGTTCGCTAAAACGC-3’ and 5’-GAACGGGTATTAAACCAAGTACCGC-3’ respectively. As we wanted a single-stranded DNA template for the mvPolX assay, asymmetric PCR (aPCR) was used to amplify the region of interest by using an unequal ratio of the primers (1:20 reverse to forward primer ratio, with 100 nM of the reverse primer) to preferentially amplify the single-stranded DNA target. All aPCR reactions were carried out with a 3 min initial denaturation at 95°C, followed by 35 cycles (denaturation at 95°C for 30 sec, annealing at 57°C for 30 sec, extension at 72°C for 15 sec) and a final extension step at 72°C for 1 min. The Thermo 2X Taq Mastermix was used for all PCR reactions carried out in the MJ mini (Bio-Rad Laboratories) thermocycler. The PCR primers were obtained from Sigma-Aldrich, India. Gel electrophoresis experiments were performed with a 3% agarose gel.

### 2.5 Spiking of environmental water sample with M13 phage

Fresh lake water was obtained from the Powai Lake in Mumbai and filtered through an 8 *µ*m filter to remove suspended particles. 100 ml aliquots of the filtered lake water were spiked with a range of concentrations of the M13 phage (from 10^3^ to 10^9^ PFU/ml) and used for genomic DNA isolation. The extracted DNA had concentrations of 16 ng/*µ*l, 14 ng/*µ*l, 19 ng/*µ*l, and 30 ng/*µ*l respectively.

### 2.6 mvPolX assay protocol

The single-stranded DNA obtained from the aPCR reaction mix binds to the probe molecule such that there is a 5’ overhang (Fig. 1C). The probe is complementary to 19 bases of the ssDNA PCR product. The detection assay (Fig. 1D) was carried out in two steps. In the first step, the two strands of the probe molecule and the target DNA strand were annealed. Initially, a probe concentration of 500 nM was used. Later we optimized it to 20 nM. The DNA fragments were added to an annealing buffer (10 mM Tris, 50 mM NaCl, pH-7.5^10^). For synthetic target sequences, the probe-to-target concentration was always maintained at 1:1. The mixture was heated at 95°C for 5 min and then cooled to 25°C over a period of 10 min. In the second step, the annealed DNA was mixed with a polymerase buffer (containing 10 mM Tris-Cl, 1 mM DTT, 0.1 mg/ml BSA, and 5% glycerol; pH 7.9^10^), and mvPolX was added to it at a final concentration of 100 nM. The entire reaction mix was incubated at 30°C for 15 min and diluted appropriately before measuring the fluorescence in a spectrofluorometer (FluoroMax-4 Spectrofluorometer, Horiba Scientific). To determine the specificity, the mvPolX assay was carried out with the 19-base single-stranded synthetic DNA target as well as a non-complementary synthetic construct. A reaction mixture without the target DNA, but with mvPolX, served as a control. To check the effect of mismatch at the 3’ end of the target DNA, two oligonucleotides with a single and a double mismatch were used. The strand displacement activity of mvPolX with our probe was visualized by polyacrylamide gel electrophoresis (PAGE) and the gel was imaged using a Amersham Typhoon V scanner (Cytiva) to see the FAM fluorescence. Along with the genomic DNA of M13 phage, we also tested a simulated environmental sample by spiking a lakewater sample with the M13 phage. All experiments were conducted in triplicates.

### 2.7 Data analysis

Unless otherwise specified, data are represented as mean and the standard error of mean (SEM) of three biological replicates. Statistical significance for the difference between samples was calculated using student’s t-test where *** indicates a p-value < 0.001 with a confidence interval of 95%.

## 3 Results and Discussion

### 3.1 Specific detection of a synthetic target sequence

We demonstrated the proof of concept of this molecular assay by designing the probe molecule for a 19-base region in gene 1 that encodes the protein G1p (or the morphogenesis protein) in the M13 phage.^13,14^ M13 is a filamentous phage with a circular single-stranded DNA as its genetic material.^15^ As M13 phage is easy to culture, we chose it for establishing the proof of our detection principle. G1p is a non-structural protein that plays a role in phage assembly ^14^ and extrusion of newly-assembled phage particles.^16^ The region of gene 1 that we selected for our probe design is conserved in M13-like filamentous phages (Fig. S1). A synthetic DNA strand (19 bases) with the same DNA sequence as the conserved region in gene 1 was used to check the specificity of the assay. A 20 bases long synthetic non-complementary strand was used as a negative control.

The polymerization and strand displacement by mvPolX were observed only in the presence of the target DNA oligonucleotide as shown in Fig. 2A (green curve). The other curves are the control reactions: (i) with only the probe and the mvPolX enzyme (red curve), and (ii) with non-complementary DNA (blue curve). Both curves show only a basal level of fluorescence. The peak fluorescence value at 520 nm from each of the curves is plotted in Fig. 2B. Compared to both negative controls, the increase in fluorescence for the target sequence showed p < 0.01, indicating it is significant.

**Figure 2.**
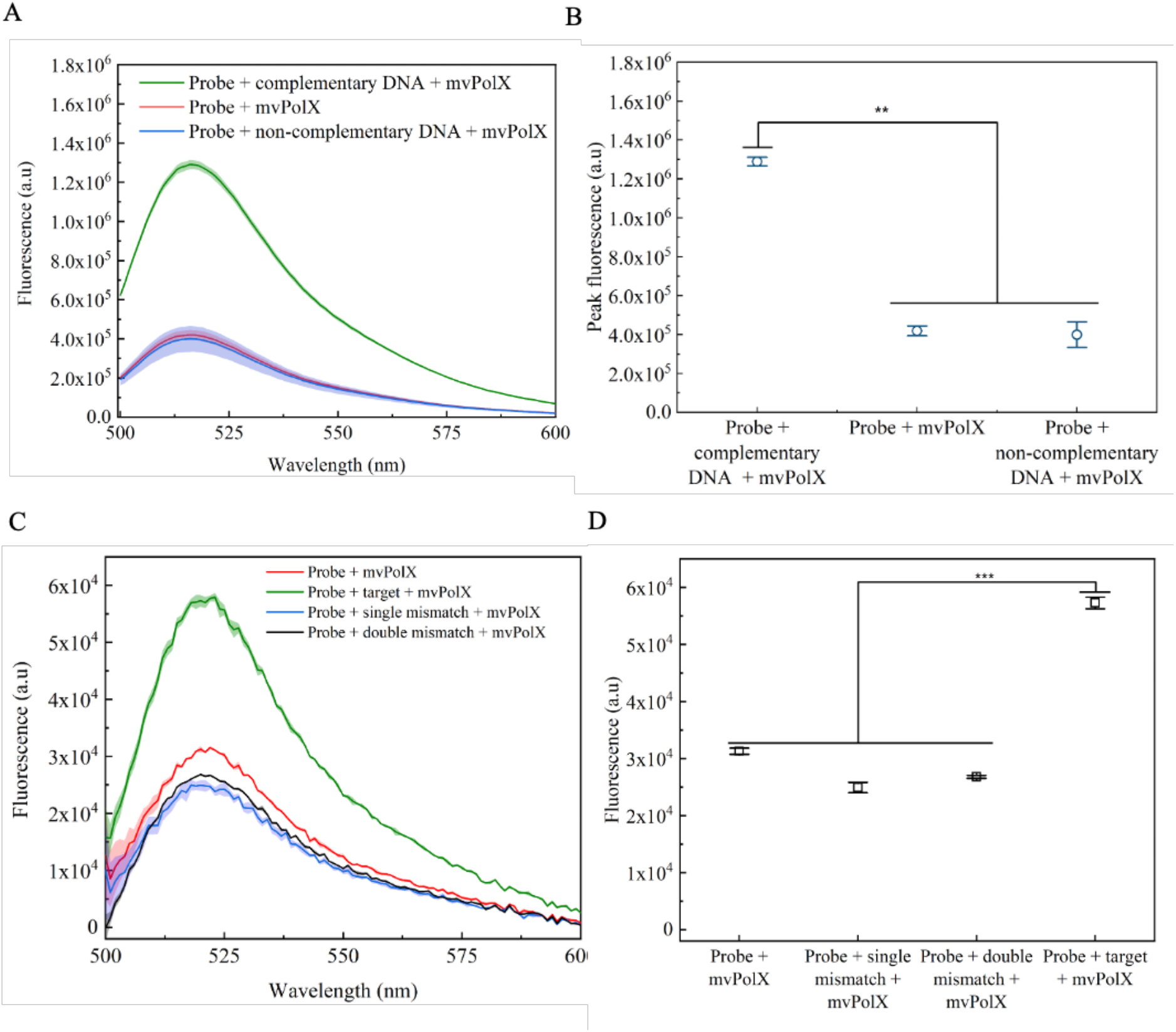
Detection of synthetic target DNA and optimization. (A) Emission spectra of the mvPolX assay with synthetic target DNA. (B) Emission maxima (520 nm) obtained from the curves in (A). (C) Emission spectra of the mvPolX assay with synthetic targets with single and double nucleotide mismatches. (D) Plot showing the emission maxima from the curves in (C). In (A) and (C), the shaded regions indicate the standard error of mean (SEM) of three biological replicates.

Next, we wished to determine how sensitive the assay is to very similar sequences. To establish the specificity, we performed the assay with synthetic targets that had a single or a double mismatch at the 3’ end. The terminal cytosine in the target sequence was replaced by adenine to introduce a single-base mismatch. To introduce the double-base mismatch, thymidine and cytosine were changed to cytosine and adenine respectively. As shown in Fig. 2C, neither polymerization nor strand displacement takes place in cases of single or double mismatch. Both these cases showed the basal level of fluorescence, which was similar to the negative control. The green curve, corresponding to the complementary target sequence, showed an increase in fluorescence. Fig. 2D shows a plot of the peak fluorescence values from each of the curves. The difference in fluorescence between the specific target and those with mismatches is statistically significant (p < 0.001). Therefore, our results indicate that this method can differentiate even a single mismatch at the 3’ end.

### 3.2 Establishing the limit of detection with the synthetic target

For determining the detection sensitivity of our assay, it is important to optimize the incubation time and the probe concentration. First, we varied the incubation time from 5 min to 30 min, while keeping the probe concentration at 500 nM for all reactions. The fluorescence signal saturates at 15 min as seen in Fig 3A. Therefore, an incubation time of 15 min at 30°C was used in all further reactions.

**Figure 3.**
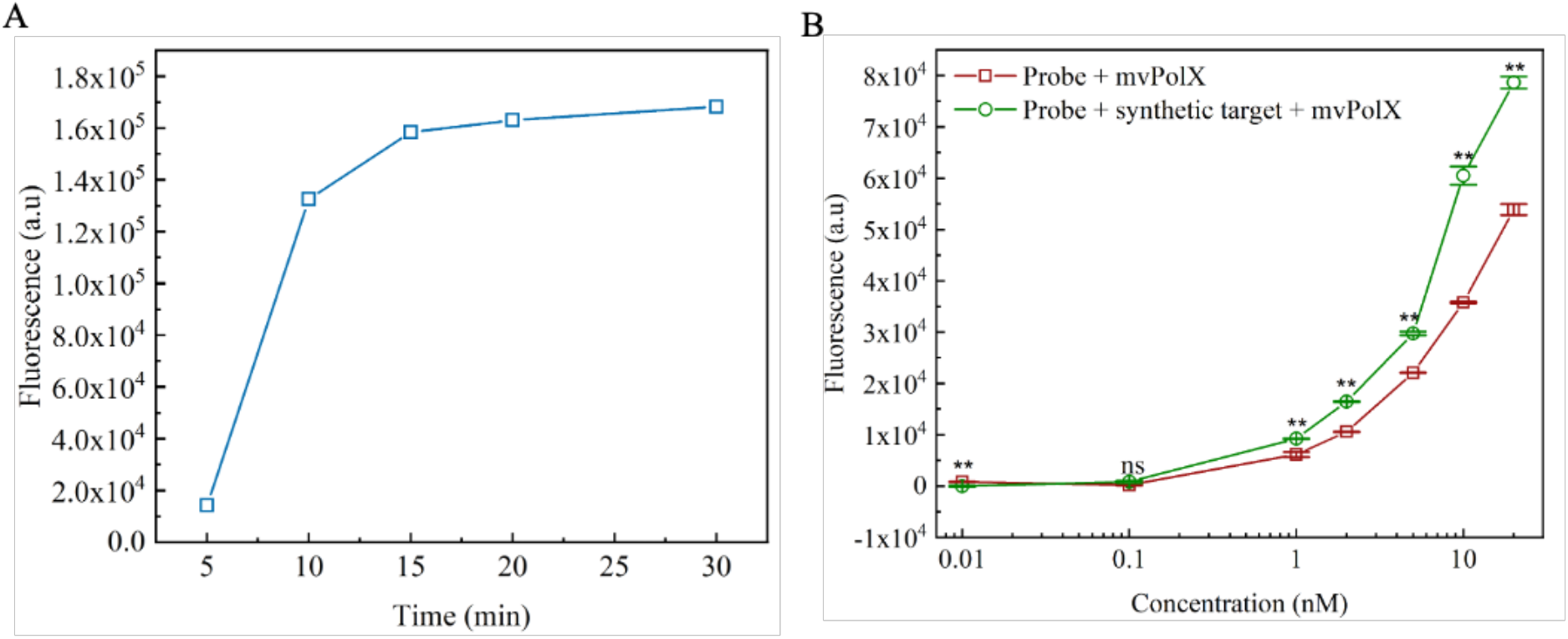
Determining the minimum reaction time and the probe concentration. (A) The fluorescence signal from the detection assay saturates at 15 min. (B) The fluorescence signal from the target can be distinguished from that of the negative control (no target) at 1 nM probe concentration.

We found that the probe itself exhibited a basal level of fluorescence. While we added equal molar concentrations of FAM- and BHQ1-labeled DNA to prepare the probe, there might be a slight difference in their actual amounts due to pipetting errors. Moreover, BHQ1 quenches only ∼93% of FAM fluorescence.^17^ The basal level of fluorescence from the probe alone can be attributed to these two reasons. Since the probe showed a basal fluorescence, it was necessary to determine the minimum probe concentration that can still allow us to separate the signal from the background. Probe concentrations ranging from 10 pM to 20 nM were tested. We observe that a minimum probe concentration of 1 nM is sufficient to detect the synthetic target DNA (p <0.01) (Fig 3B).

The limit of detection (LOD) was calculated from a calibration curve (Fig. S2) of the fluorescence readings for different concentrations of the substrate (i.e. target-bound probe). The synthetic M13 target DNA was used for generating the calibration curve. The LOD was calculated using the formula, 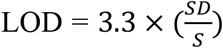, where SD is the standard deviation and S is the slope of the calibration curve. Using this formula, the LOD of our assay was found to be 650 pM.

### 3.3 Detection of 10 copies of genomic DNA

After establishing the sensitivity and specificity of our assay with a 19-base synthetic DNA target, we wanted to validate it with a longer sequence of genomic DNA of M13 phage. Since this method requires the target DNA to have a 3’OH group that orients to bind adjacent to the gap region, we needed to perform 35 cycles of asymmetric PCR (aPCR) to generate the target ssDNA. Fig. 4A shows the gel electrophoresis image of the aPCR product in lane A and the no template control (NTC) in Lane B. The prominent band at 158 bp in lane A corresponds to the dsDNA obtained from aPCR, while the faint band just below corresponds to the ssDNA product. Fig. 4B shows the fluorescence emission spectra after the mvPolX assay. The test reaction, with the ssDNA target and mvPolX (green curve), shows a sizable increase in fluorescence compared to the various negative controls. The peak fluorescence values (at 520 nm) from the reaction and the negative controls are plotted in Fig. 4C. As seen from the plot, the difference in fluorescence between the reaction and the negative controls are significant (p < 0.05). Fig. 4D shows the peak fluorescence values for different copy numbers of template DNA. We can detect as low as 10 copy numbers (p < 0.05). Further, as shown in Fig. S3 of Supplementary Information, we also confirmed that the increase in fluorescence is primarily due to the ssDNA generated by aPCR.

**Figure 4.**
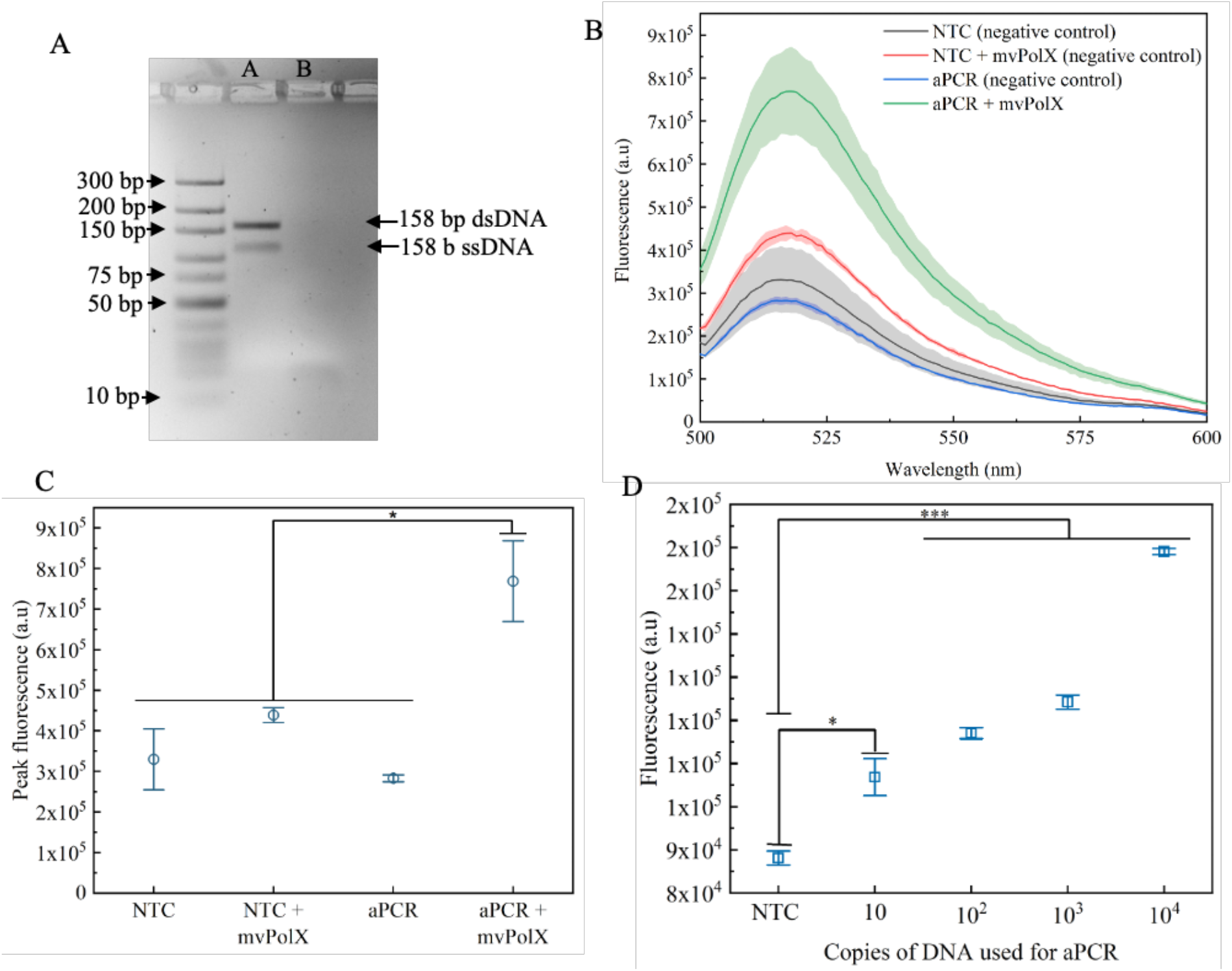
Detection of target using genomic DNA. (A) Agarose gel electrophoresis of the asymmetric PCR product. Lane A shows both the double-stranded (between 150 and 200 bp bands) and the single-stranded PCR products (below the 150 bp band). Lane B is the no-template control (NTC). (B) Fluorescence emission spectra showing an increase in fluorescence in presence of the target and mvPolX. The other curves are the negative controls. The shaded regions indicate the SEM from three biological replicates. (C) Scatter plot of the peak fluorescence emission (520 nm) from (B). The increase in fluorescence is statistically significant compared to the negative controls. (D) Plot showing the peak fluorescence emission (520 nm) from the mvPolX assay with different copy numbers.

### 3.4 Confirmation of strand displacement activity of mvPolX with our probe

We performed PAGE after the mvPolX assay to confirm its strand displacement activity with respect to our probe. ssDNA migrates faster than both dsDNA or partially dsDNA of the same size. As shown in Fig. 5, except lanes D and H, all other lanes are negative controls. Lane D, containing the synthetic target, shows a stronger FAM fluorescence compared to the respective negative controls (lanes A – C).

**Figure 5.**
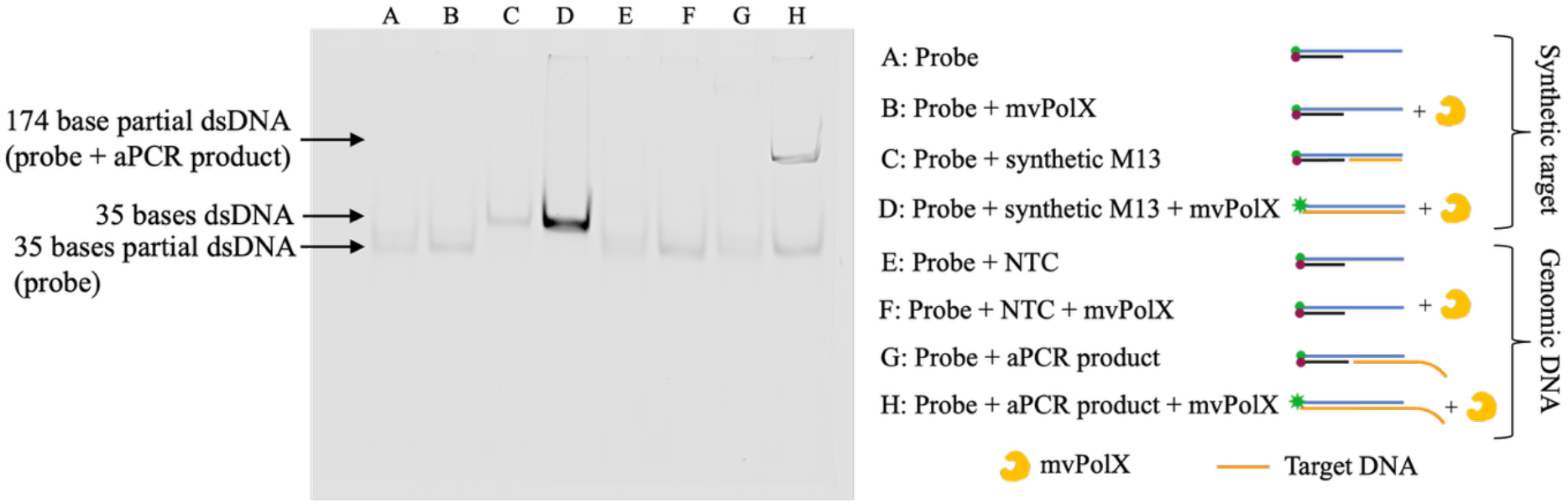
FAM fluorescence from the image of a PAGE gel. Lanes A - D correspond to the synthetic target sequence and lanes E - H correspond to the genomic DNA from the M13 phage. Lane D has a stronger FAM fluorescence compared to its negative controls (A - C). Only lane H shows the presence of the 158-base genomic DNA fragment unlike its negative controls (lanes E – G).

Lane H corresponds to the test reaction with the aPCR product annealed to the probe and mvPolX. The band with the higher molecular weight (158 bases) is seen only at lane H and corresponds to the FAM fluorescence from the target after the BHQ1 strand is displaced. Therefore, our results confirm that the strand displacement activity of mvPolX takes place in the molecular assay that we have designed.

### 3.5 Detection of target DNA in a lakewater environment

Finally, we wanted to see if the assay works in an environmental sample. Therefore, we spiked a lakewater sample with the M13 phage and performed the mvPolX assay. Fig. 6A shows a schematic representation of the entire process. We prepared lakewater samples by spiking 10^3^ PFU/ml, 10^5^ PFU/ml, 10^7^ PFU/ml and 10^9^ PFU/ml of the phage suspension. Fig. 6B shows the fluorescence curves for different phage concentrations, while Fig. 6C shows the corresponding peak fluorescence values (520 nm). We could detect DNA from 10^5^ PFU/ml onwards (p < 0.001). We could not distinguish between the NTC and the sample spiked with 10^3^ PFU/ml, indicating that the detection technique needs to be improved further for handling more complex samples. This loss of sensitivity may possibly be due to inefficiency of the phage capture and/or the DNA extraction process. In spite of the reduced sensitivity, our results show that the assay can work, in principle, in real samples.

**Figure 6.**
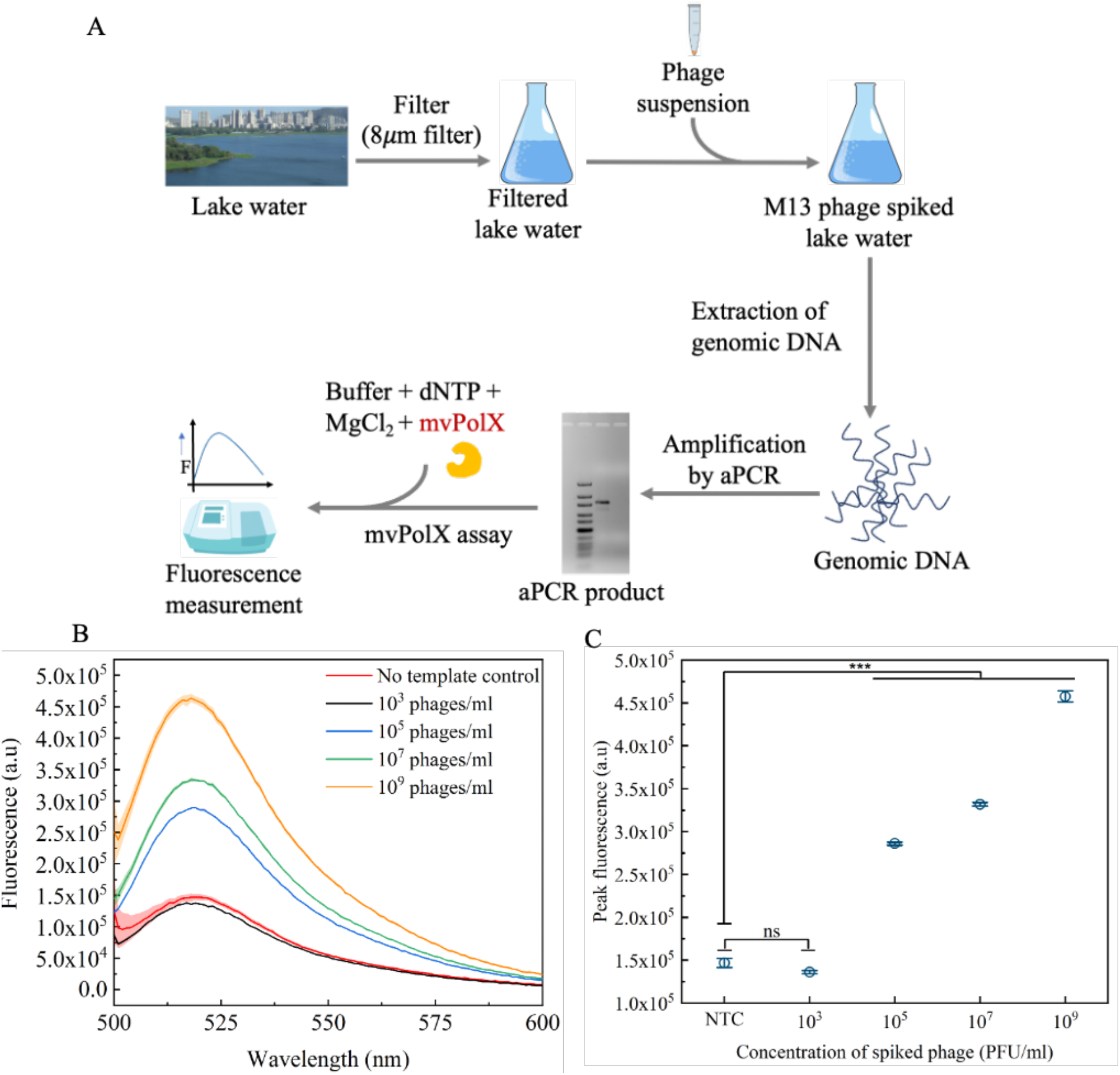
Detection of target DNA in a lakewater environment. (A) Schematic work flow for performing the mvPolX assay on a lakewater sample spiked with M13 phage. (B) Plot showing the fluorescence emission for different phage concentrations. The shaded regions indicate SEM from three technical replicates. (C) Plot showing the peak fluorescence values (520 nm) obtained from (B).

## Conclusion

Our molecular detection method, using the mvPolX enzyme, demonstrates detection of target DNA using a suitably designed probe. It requires only 15 min of incubation with mvPolX at 30°C. The method is highly specific and can distinguish between single-base mismatches at the 3’ end. It has a LOD of 650 pM when used to detect a synthetic target. When combined with an optimized aPCR protocol, this method can detect 10 copies of the target DNA. We demonstrated this assay with M13 phages spiked in an environmental sample. In summary, our molecular assay is a rapid, specific and sensitive method for detecting target DNA. While we establish the proof of principle with a phage here, it has the potential to be extended for SNP detection. Using probes coupled with different fluorophores to detect multiple DNA target regions, this can be developed into a full-fledged multiplexed diagnostic platform.

## Acknowledgements

The authors thank Wadhwani Research Centre for Bioengineering (WRCB) (DO/2022-WRCB002-082) for partial funding support towards consumables. They also thank Science and Engineering Research Board, SERB (CRG/2023/001315), the Department of Biotechnology, DBT [BT/PR35928/BRB/10/1841/2019], and The Board of Research in Nuclear Sciences, BRNS [58/14/11/2020-BRNS/37188] grants for funding support. The authors thank the common equipment facility of the Department of Biosciences and Bioengineering, IIT Bombay, for the use of gel documentation system and the multimodal plate reader. The authors acknowledge Prof. Santanu Kumar Ghosh for providing the *E. coli* XL1-Blue strain and Prof. Ashutosh Kumar for providing access to the FluoroMax-4 Spectrofluorometer. The authors thank and acknowledge Department of Chemistry, IIT Bombay, for access to the Amersham Typhoon V imager.

