## Supplementary Data for "Strand Displacement Activity of Mimiviral Polymerase X Enables Rapid Detection of Sequence-Specific DNA Targets"

### Supporting Information

#### 1. Alignment showing the conservation of the chosen sequence in gene 1 of M13-like phages

To identify the conserved region in Gene 1, 21 sequences of M13 like filamentous phages (f1 and fd phages) were retrieved from NCBI database. These sequences were aligned using multiple alignment using fast fourier transform (MAFFT). Fig S1 shows the MAFFT view of a 19 base pair conserved region that was identified and the single stranded region of the probe was designed complementary to this region.

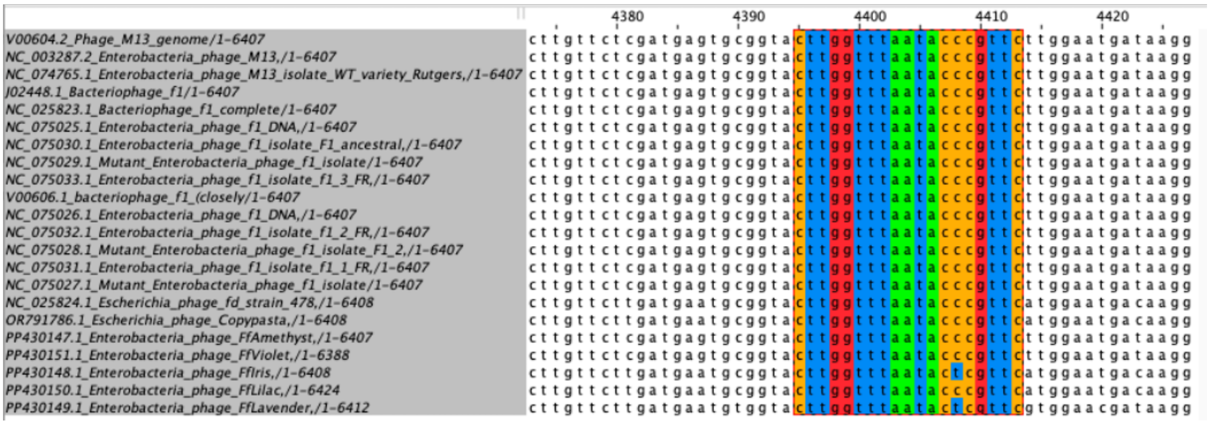


**Figure S1**. Alignment of gene 1 in all M13 like phages showing region of conservation

#### 2. Generation of the calibration curve for the synthetic target sequence

A calibration curve was generated by plotting the fluorescence measurements of mvPolX against the concentration. Various concentration of probe (0.01 nM to 10 nM) with a 1:1 molar ratio of target for each probe concentration were used to generate this plot. The linear fit equation is $y = 5888.6x + 1685.8$

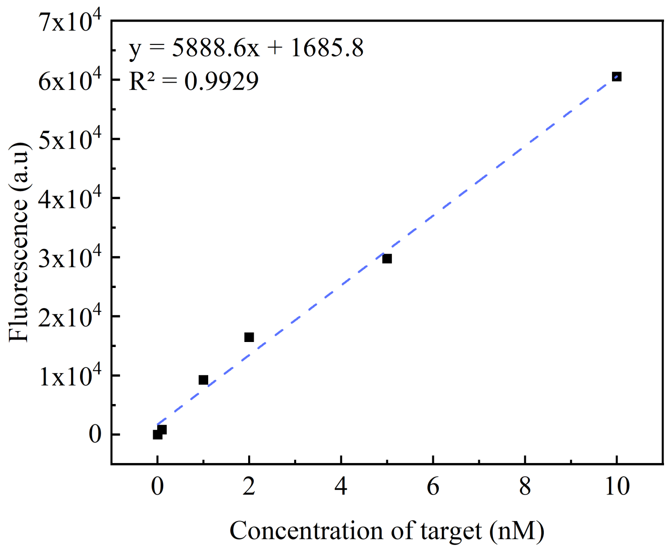


**Figure S2.** Calibration curve for determination of LOD using the synthetic target sequence.

### 3. The increase in fluorescence is due to ssDNA target

We confirmed that the increase in fluorescence is primarily due to ssDNA generated during aPCR. To do that, we performed aPCR for 5 to 30 cycles, and plotted the resulting fluorescence intensity (Fig. S3A). As expected, there was an increase in fluorescence with the increasing number of aPCR cycles. To determine the relative amounts of ssDNA and dsDNA produced during different cycles of aPCR, we ran the samples on an agarose gel (Fig S3B) and used ImageJ to analyze the band intensity (Fig. S3C). We found that the intensity of dsDNA saturates by the $10^{th}$ cycle, while the intensities of the ssDNA bands kept on increasing with the cycle number. This observation confirms that the increase in the fluorescence intensity in the mvPolX assay is due to ssDNA and not dsDNA. Further, we calculated by which cycle the reverse primer gets exhausted, leading to formation of ssDNA. We started aPCR with $10^9$ copies of genomic DNA, while the number of copies of reverse primer was $1.2 \times 10^{12}$. Using these values, we calculated that the reverse primer gets exhausted by the ~$7^{th}$ cycle of aPCR, which roughly matches with our experimental conclusions.

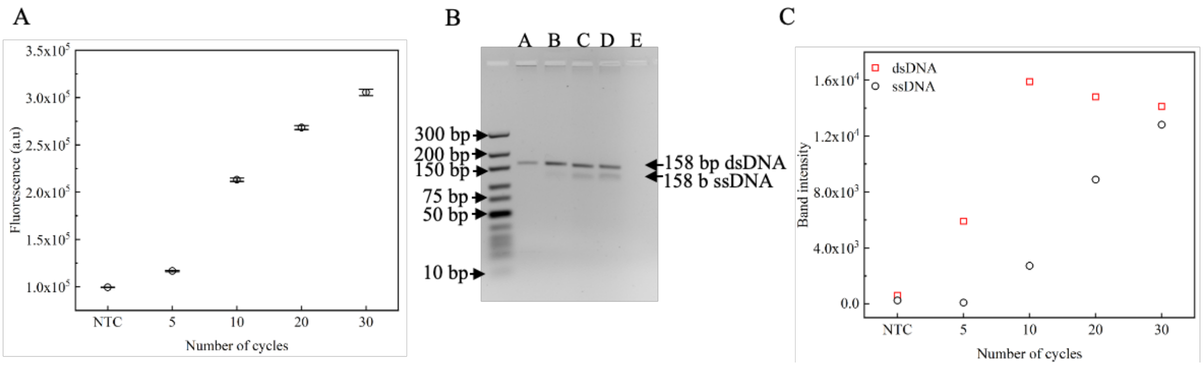


**Figure S3.** (A) Plot of the peak fluorescence emission at 520 nm for different cycle numbers of aPCR. (B) Agarose gel image showing the aPCR products after 5, 10, 20 and 30 cycles. (C) Plot showing the intensities of the dsDNA and ssDNA bands in agarose gel.
